# Differential Proteomic Analysis of Serum and Bone Marrow Supernatant in Acute myeloid leukemia Patients at Low and Intermediate Altitudes after Chemotherapy

**DOI:** 10.1101/2024.04.09.588705

**Authors:** Qi Sun, Houfa Zhou, Aibo Wang, Wenqian Li, Youbang Xie

## Abstract

Chemotherapy is the primary treatment for patients with acute myeloid leukemia (AML). In addition to factors such as patient age, physical condition, and choice of medication, we have noticed that environmental factors such as altitude may also have a significant impact on post-chemotherapy bone marrow suppression in AML patients in clinical practice. The results indicate that there are differences in the proteomics of the two groups of patients during the bone marrow suppression period after chemotherapy. Differentially expressed proteins are primarily located in the cytoplasm, extracellular space, and nucleus, followed by mitochondria and membranes. These differentially expressed proteins mainly participate in biological processes such as cell and metabolism. For differential protein KEGG pathway enrichment analysis, it was found that metabolic pathways were mainly enriched in the metabolic category, while the PI3K-Akt signaling pathway, HIF-1 signaling pathway, NF-κB signaling pathway, and calcium signaling pathway were enriched in the signaling pathways.

## Introduction

Acute myeloid leukemia (AML) is a malignant clonal disease of hematopoietic stem and progenitor cells. Chemotherapy, as a significant treatment modality for AML patients, while effective, also induces certain toxic side effects, with bone marrow suppression being the most common. According to WHO statistics in 2020, there were 19.29 million new cancer cases worldwide, with 80% of patients experiencing bone marrow suppression during tumor chemotherapy or radiotherapy. This suppression often leads to dose reduction, treatment limitations, or even discontinuation, which are significant obstacles in cancer treatment(Nian *et al*, 2016). Apart from patient age, physical condition, and drug selection, environmental factors such as altitude may also exert significant influence on post-chemotherapy bone marrow suppression in AML patients. Following standard-dose chemotherapy, AML patients at moderate altitudes experience significantly exacerbated bone marrow suppression, with prolonged recovery times for bone marrow hematopoietic function, lasting up to approximately 4-5 weeks in some cases. Due to significant differences in the living environment and lifestyle of patients residing at moderate altitudes compared to those at low altitudes, coupled with variations in climatic conditions, atmospheric pressure, and oxygen concentration associated with altitude, moderate altitude regions may exert a notable influence on the physiological status of individuals. Proteins, as high-throughput biological entities, directly influence biological physiological functions and pathological processes through processes such as transcription, translation, and post-translational modifications(Li *et al*, 2017). Proteomics, as an effective tool for studying protein expression and functionality, has demonstrated immense potential in diseases such as cancer. However, there is currently a lack of in-depth research regarding the differential effects of post-chemotherapy bone marrow suppression among AML patients residing at different altitudes.Understanding the impact of altitude factors on bone marrow suppression, particularly at the protein level, will contribute to a better comprehension of treatment responses and prognoses among AML patients in diverse environmental settings.

## Results

### Baseline characteristics

In the low-altitude group, there were 5 cases with an average age of (42.00±20.67) years, including 3 females (60.0%) and 2 males (40.0%). In the mid-altitude group, there were 5 cases with an average age of (51.80±17.11) years, including 1 female (20.0%) and 4 males (80.0%). Among them, in the low-altitude group, there were 2 cases of M2 type, 2 cases of M4 type, and 1 case of M5 type, while in the mid-altitude group, there were 1 case of M2 type, 2 cases of M4 type, and 2 cases of M5 type. There were no statistically significant differences in age, gender, or FAB classification between the groups (*P* > 0.05), and the baselines were comparable (see Tables 1 and 2). The difference in bone marrow suppression degree between the two groups at 8 days of chemotherapy was not statistically significant (*Z* = -1.000, *P* = 0.317). However, the difference in bone marrow suppression degree between the two groups at 14 days and 28 days of chemotherapy was statistically significant (*Z* = -1.936, *P* = 0.053; *Z* = -2.169, *P* = 0.030) (see Figure 1).

**Table 1:**
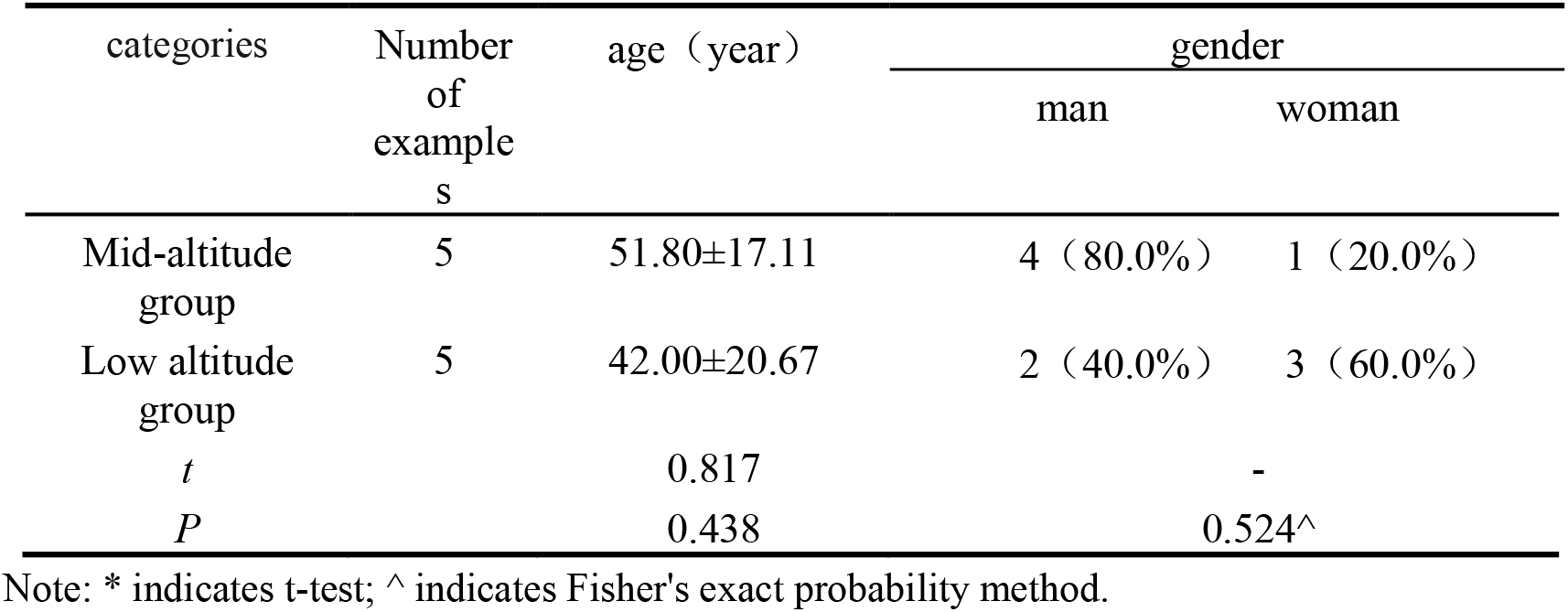
Comparison of Age and Gender between Low and Mid-Altitude Groups 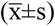 [n (%)]

| categories | Number of examples | age (year) | gender |  |
| --- | --- | --- | --- | --- |
|  |  |  | man | woman |
| Mid-altitude group | 5 | 51.80±17.11 | 4 (80.0%) | 1 (20.0%) |
| Low altitude group | 5 | 42.00±20.67 | 2 (40.0%) | 3 (60.0%) |
| *t* |  | 0.817 |  | - |
| *P* |  | 0.438 |  | 0.524^ |

Note: \* indicates t-test; ^ indicates Fisher's exact probability method.

**Table 2:**
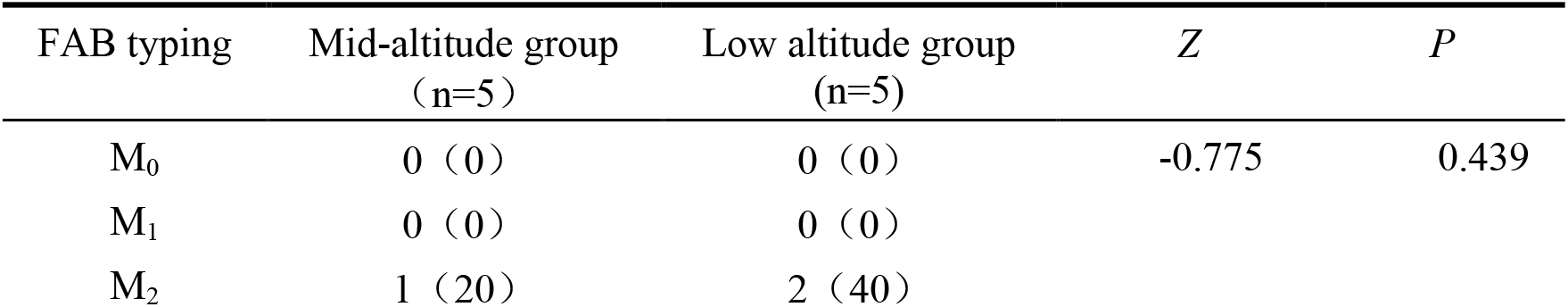
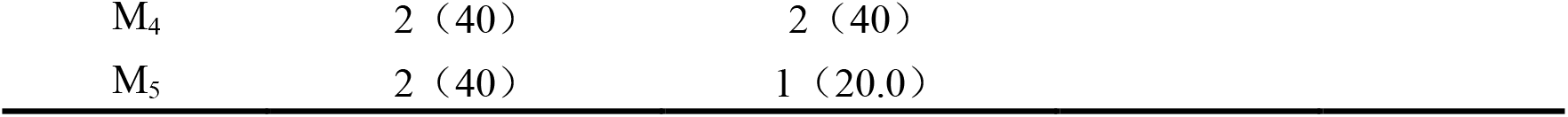
Comparison of Subtypes between Low-Altitude Group and Mid-Altitude Group [n (%)]

| FAB typing | Mid-altitude group (n=5) | Low altitude group (n=5) | *Z* | *P* |
| --- | --- | --- | --- | --- |
| M<sub>0</sub> | 0 (0) | 0 (0) | -0.775 | 0.439 |
| M<sub>1</sub> | 0 (0) | 0 (0) |  |  |
| M<sub>2</sub> | 1 (20) | 2 (40) |  |  |

|  |  |  |
| --- | --- | --- |
| M<sub>4</sub> | 2 (40) | 2 (40) |
| M<sub>5</sub> | 2 (40) | 1 (20.0) |

**Figure 1:**
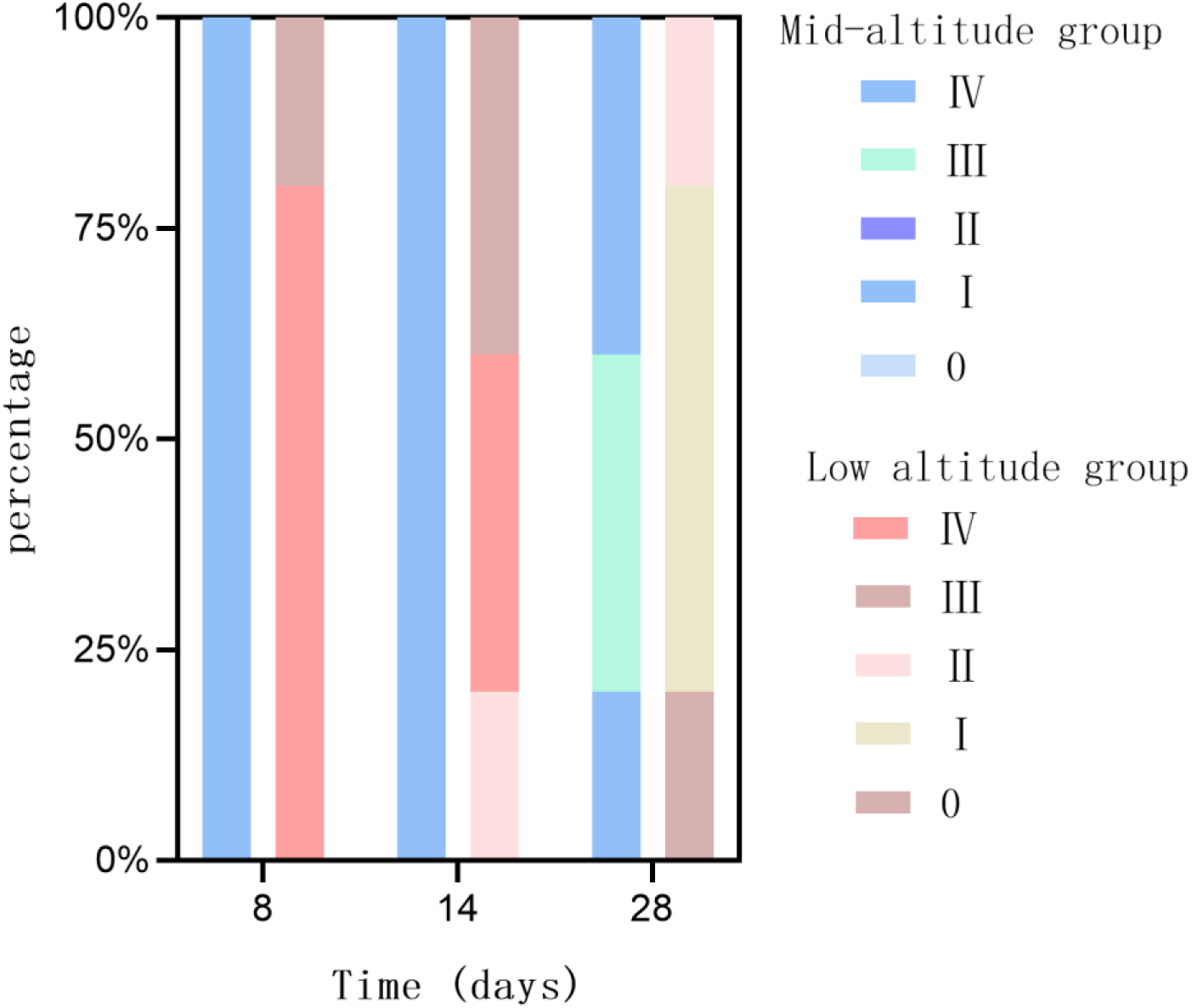
Stacked plot comparing myelosuppressive indexes at low and middle altitudes.

### Screening of differentially expressed proteins

In the analysis of differential proteins, proteins with FC (Fold Change) > 1.5 or < 0.6667 and *P* < 0.05 were defined as significantly different proteins. Specifically, when *P* < 0.05, FC > 1.5 was considered significantly upregulated, and FC < 0.6667 was considered significantly downregulated. In peripheral serum at 8 days after chemotherapy, compared to the low-altitude AML group, there were 78 different proteins in the mid-altitude AML group, including 52 upregulated proteins and 26 downregulated proteins. In peripheral serum at 28 days after chemotherapy, compared to the low-altitude AML group, there were 86 different proteins in the mid-altitude AML group, including 59 upregulated proteins and 27 downregulated proteins. In bone marrow supernatant at 28 days after chemotherapy, compared to the low-altitude AML group, there were 43 different proteins in the mid-altitude AML group, including 32 upregulated proteins and 11 downregulated proteins. The protein quantification results were displayed in volcano plots, as shown in Figure 2.

**Figure 2:**
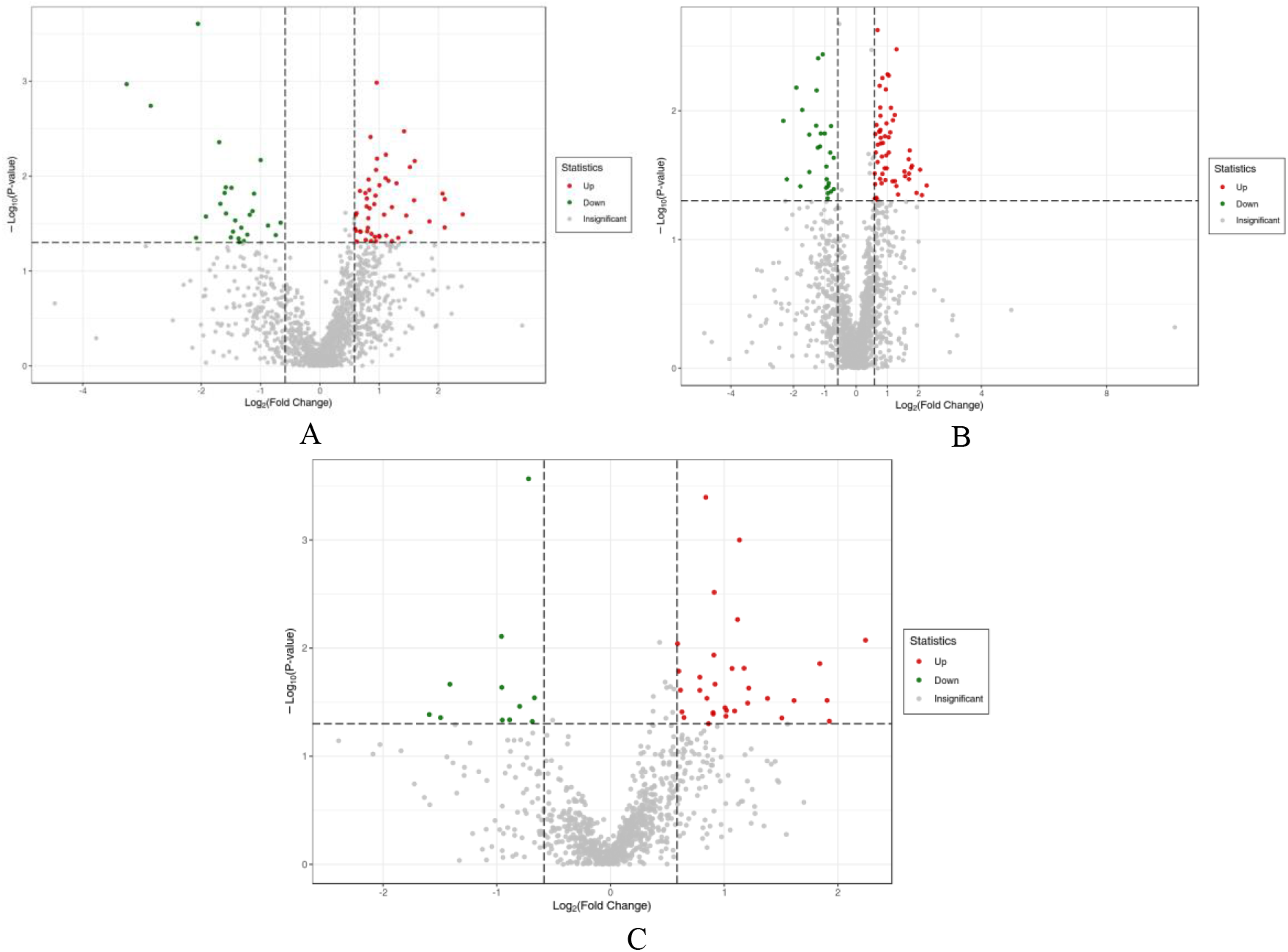
Volcano plots illustrating differential expression protein quantification comparisons between low-altitude and mid-altitude groups in peripheral serum at 8 days after chemotherapy, peripheral serum at 28 days after chemotherapy, and bone marrow supernatant. A. Peripheral serum at 8 days after chemotherapy: Low-altitude group vs. Mid-altitude group up-regulated: 52, down-regulated: 26 B. Peripheral serum at 28 days after chemotherapy: Low-altitude group vs. Mid-altitude group up-regulated: 59, down-regulated: 27 C. Bone marrow supernatant at 28 days after chemotherapy: Low-altitude group vs. Mid-altitude group up-regulated: 32, down-regulated: 11 The x-axis represents the log2 fold change, and the y-axis represents -log10P value. The red and green dots represent upregulated and downregulated proteins, respectively.

**Figure 3:**
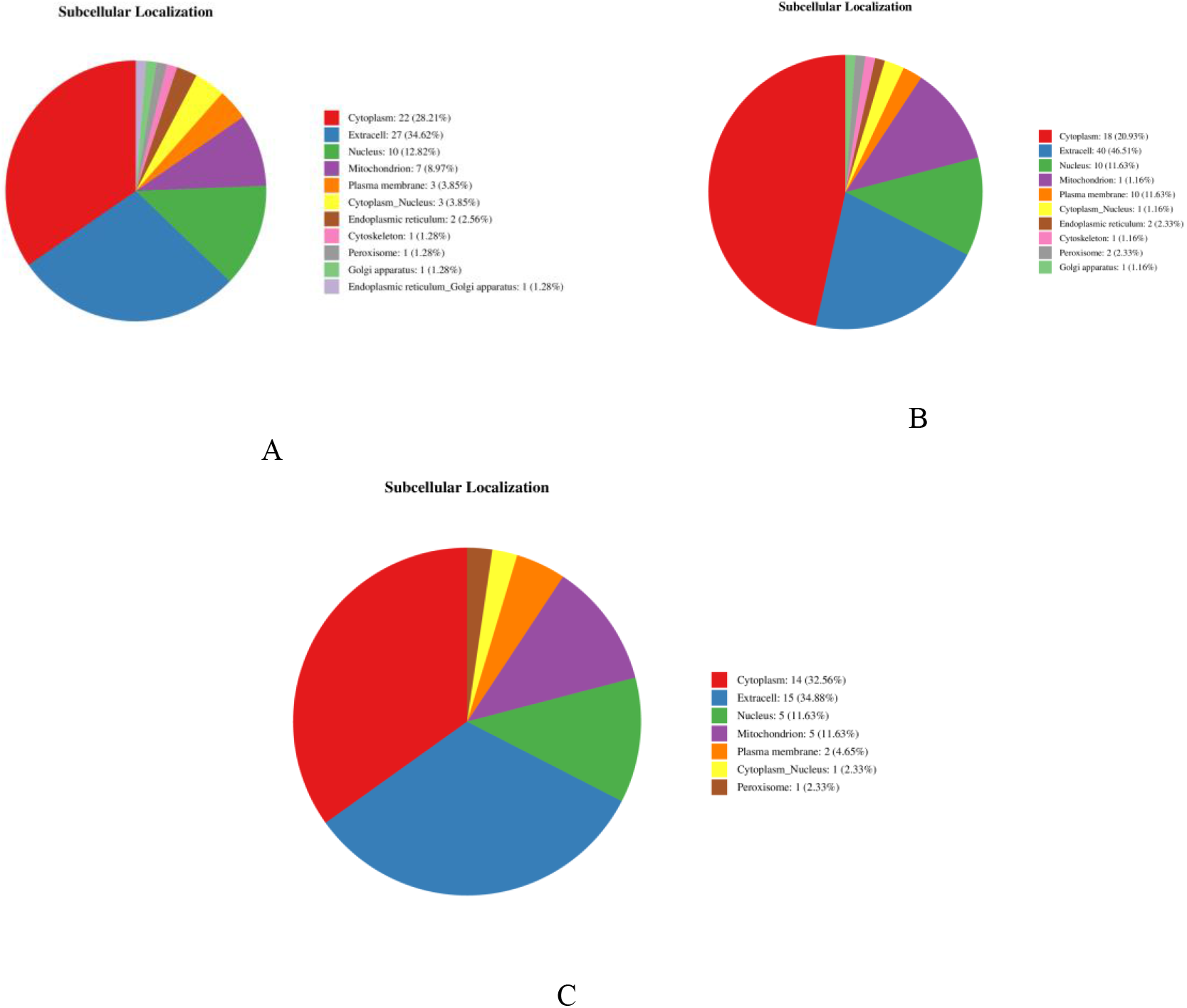
Subcellular Localization Distribution of Differentially Expressed Proteins. Note: A and B represent the subcellular localization distribution of differentially expressed proteins in the low-altitude group and the mid-altitude group at 8 and 28 days after chemotherapy in peripheral blood serum, respectively. C represents the subcellular localization distribution of differentially expressed proteins in the low-altitude group and the mid-altitude group at 28 days after chemotherapy in bone marrow supernatant. The numbers in the pie charts represent the quantity of significantly differentially expressed proteins, while the percentages represent the proportion of differentially expressed proteins in the total.

**Figure 4:**
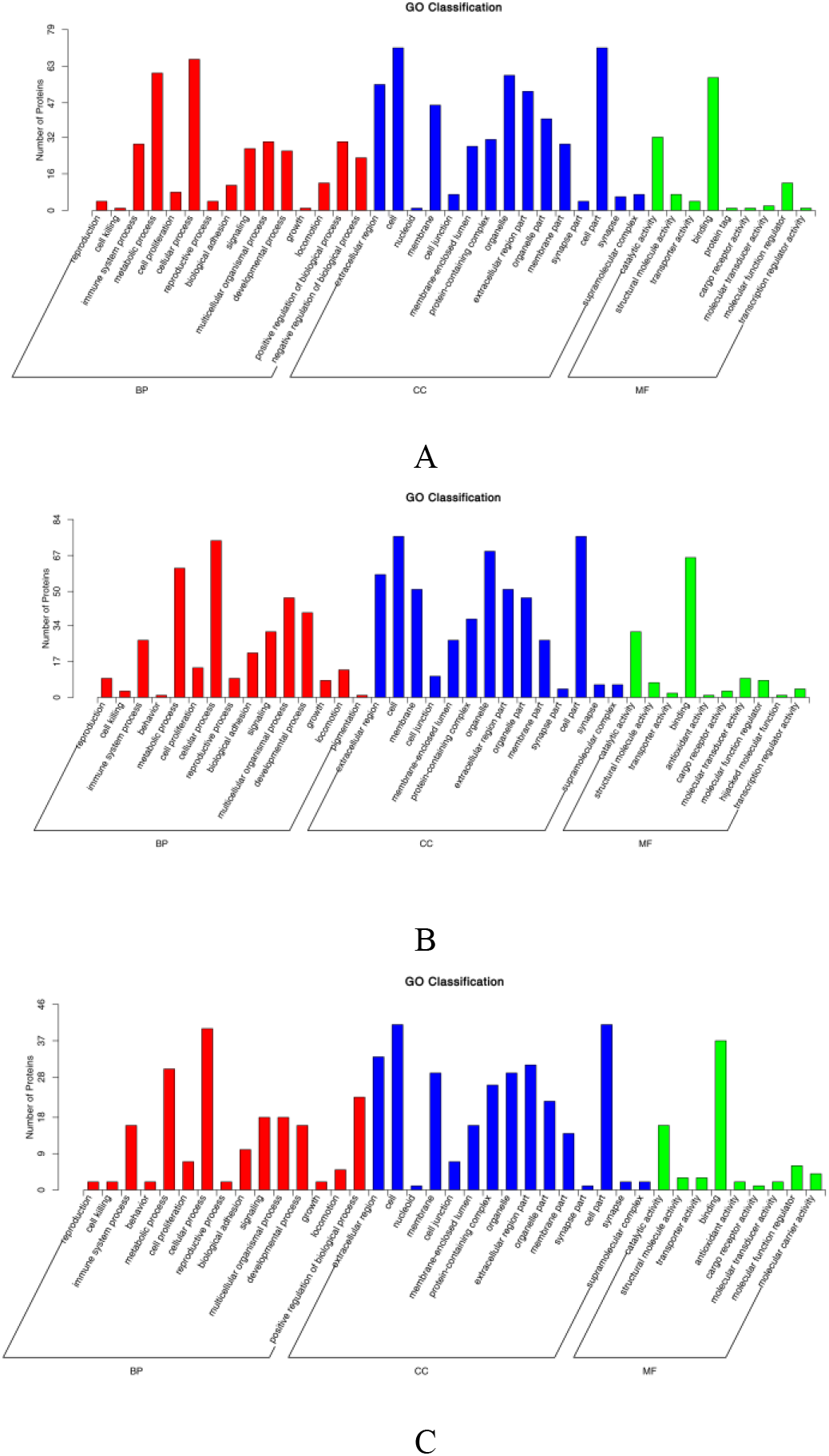
Bar Chart of GO Classification. Note: A and B respectively represent the statistical distribution of differentially expressed proteins in the GO secondary classification of peripheral blood serum after 8d and 28d of chemotherapy in low-altitude and mid-altitude groups; C represents the statistical distribution of differentially expressed proteins in the GO secondary classification of bone marrow supernatant after 28d of chemotherapy in low-altitude and mid-altitude groups. The x-axis represents the secondary GO entries, the y-axis represents the number of differentially expressed proteins for each GO entry, and the different colors of the bars represent different primary classifications.

### Subcellular localization of differential proteins

Using the WoLF PSORT software, the subcellular localization analysis of differentially expressed proteins between the two groups revealed the following results (Figure).

For the differential proteins in peripheral serum at 8 days after chemotherapy, subcellular localization was predominantly in the Cytoplasm, Extracell, and Nucleus, followed by Mitochondron and Plasma membrane, Cytoplasm-Nucleus, Endoplasmic reticulum. Additionally, four differential proteins were localized in the Cytoskeletion, Peroxisome, Golgi apparatus, and Endoplasmic reticulum-Golgi apparatus.

For the differential proteins in peripheral serum at 28 days after chemotherapy, subcellular localization was predominantly in the Cytoplasm, Extracell, and Nucleus, followed by Mitochondria and Plasma membranes, Cytoplasm-Nucleus, Endoplasmic reticulum, and Cytoskeleton. Additionally, two differential proteins were localized in the Peroxisome and Golgi apparatus.

For the differential proteins in bone marrow supernatant at 28 days after chemotherapy, subcellular localization was predominantly in the Cytoplasm, Extracell, and Nucleus, followed by Mitochondria and Plasma membrane. Additionally, two differential proteins were localized in the Cytoplasm-nucleus and Peroxisome.

### GO functional annotation of differential proteins

Gene Ontology (GO), available at http://geneontology.org/, is an internationally standardized classification system for gene functions, used to describe various attributes of genes and gene products. GO annotations mainly consist of three parts: Molecular Function (MF), Biological Process (BP), and Cellular Component (CC). The number of proteins corresponding to GO nodes in BP, CC, and MF is listed, and the distribution of differentially expressed proteins in the second-level GO annotation is statistically analyzed (Figure 5).

**Figure 5:**
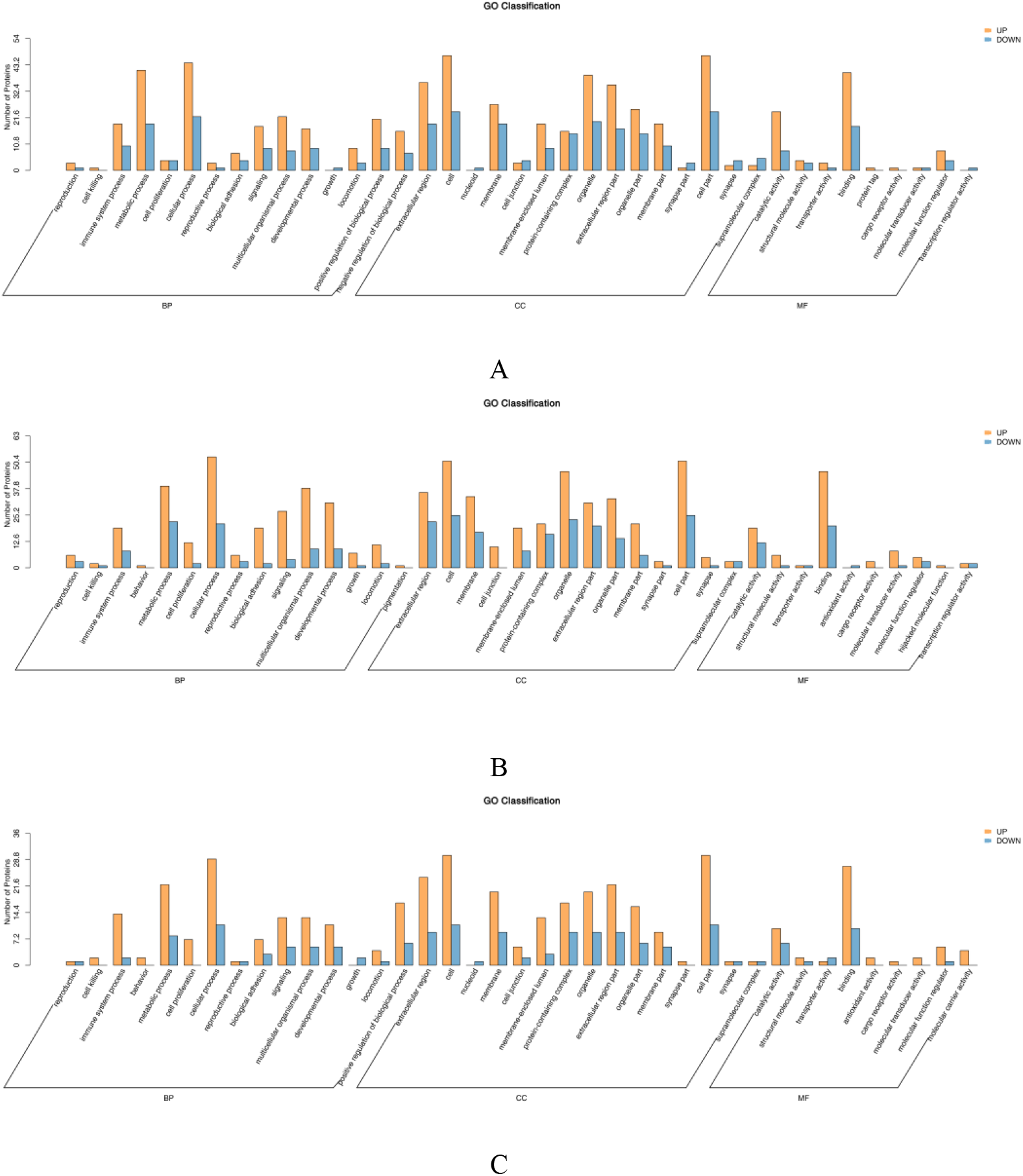
Bar Chart of GO Classification for Up- and Down-Regulated Proteins. Note: A and B respectively represent the statistical distribution of up-regulated and down-regulated proteins in the peripheral serum of patients in the low-altitude group and the mid-altitude group at 8d and 28d after chemotherapy in the second-level GO classification; C represents the statistical distribution of up-regulated and down-regulated proteins in the bone marrow supernatant of the low-altitude group and the mid-altitude group at 28d after chemotherapy in the second-level GO classification. The horizontal axis represents the second-level GO terms, and the vertical axis represents the number of differentially expressed proteins for each GO term. Yellow indicates up-regulated proteins, while blue indicates down-regulated proteins.

In the category of Biological Processes, the proportion of differentially expressed proteins between the low-altitude AML group and the mid-altitude AML group in peripheral blood serum at 8 days and 28 days after chemotherapy, as well as in bone marrow supernatant at 28 days after chemotherapy, is mostly related to cellular process, followed by metabolic processes. In the category of Cellular Components, the most abundant component is the cell, followed by the cell part. Regarding Molecular Function, the majority of differentially expressed proteins are involved in both binding and catalytic activity.

Among all subcategories, the largest number of differentially expressed proteins is annotated with terms related to cellular processes, metabolic processes, cell, cell part, binding, catalytic activity, and binding annotation. At the same time, the top 50 GO-Terms with enriched analysis results (ranked from smallest to largest P-value) were selected to create an enrichment entry bar graph (Figure 6). It was observed that the differentially expressed proteins in peripheral blood serum at 8 days and 28 days after chemotherapy, as well as in bone marrow supernatant at 28 days after chemotherapy, mainly participate in biological processes. Specifically, in peripheral blood serum at 8 days after chemotherapy, differentially expressed proteins are mainly involved in responses to biological stimuli, responses to other organisms, and responses to external biological stimuli. In peripheral blood serum at 28 days after chemotherapy, differentially expressed proteins mainly participate in participate in biological adhesion, cell adhesion, and positive regulation of the immune system process. In bone marrow supernatant at 28 days after chemotherapy, differentially expressed proteins mainly responses to other organisms, responses to external biological stimuli, and responses to bacterium.

**Figure 6:**
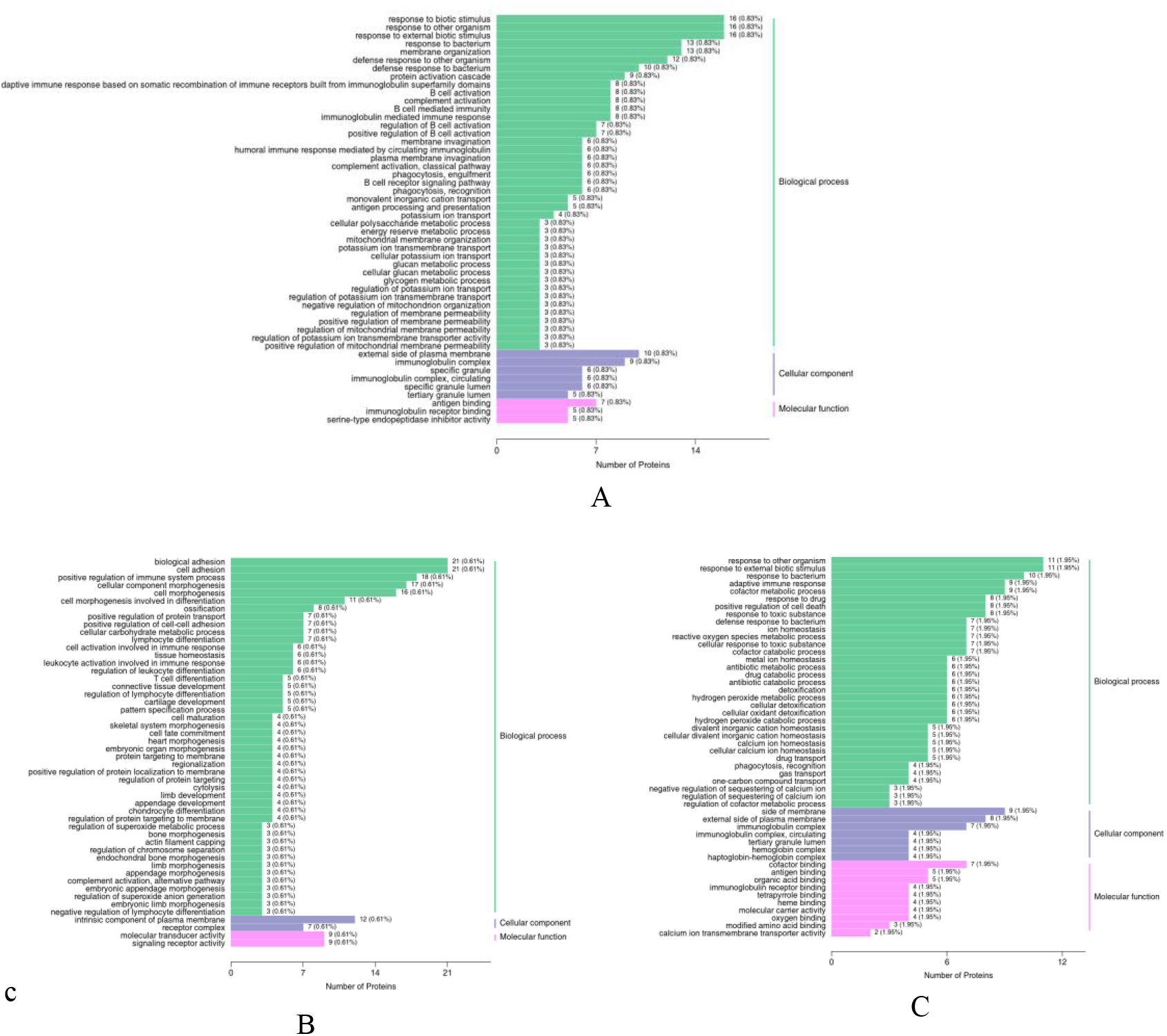
Bar graph of GO functional enrichment analysis of differential proteins. Note: A and B respectively represent the enrichment analysis of differential proteins in BP, CC, and MF categories of peripheral serum samples from the low-altitude group and the mid-altitude group at 8d and 28d after chemotherapy; C represents the enrichment analysis of differential proteins in BP, CC, and MF categories of bone marrow supernatant samples from the low-altitude group and the mid-altitude group at 28d after chemotherapy. The horizontal axis represents the annotated differential proteins, and the vertical axis represents the names of GO terms. The numbers in the graph indicate the number of annotated differential proteins for each GO term, with the specific value of the ratio of the number of annotated differential proteins to the total number of annotated differential proteins in parentheses. The label on the far right represents the primary classification to which the GO term belongs.

**Figure 7:**
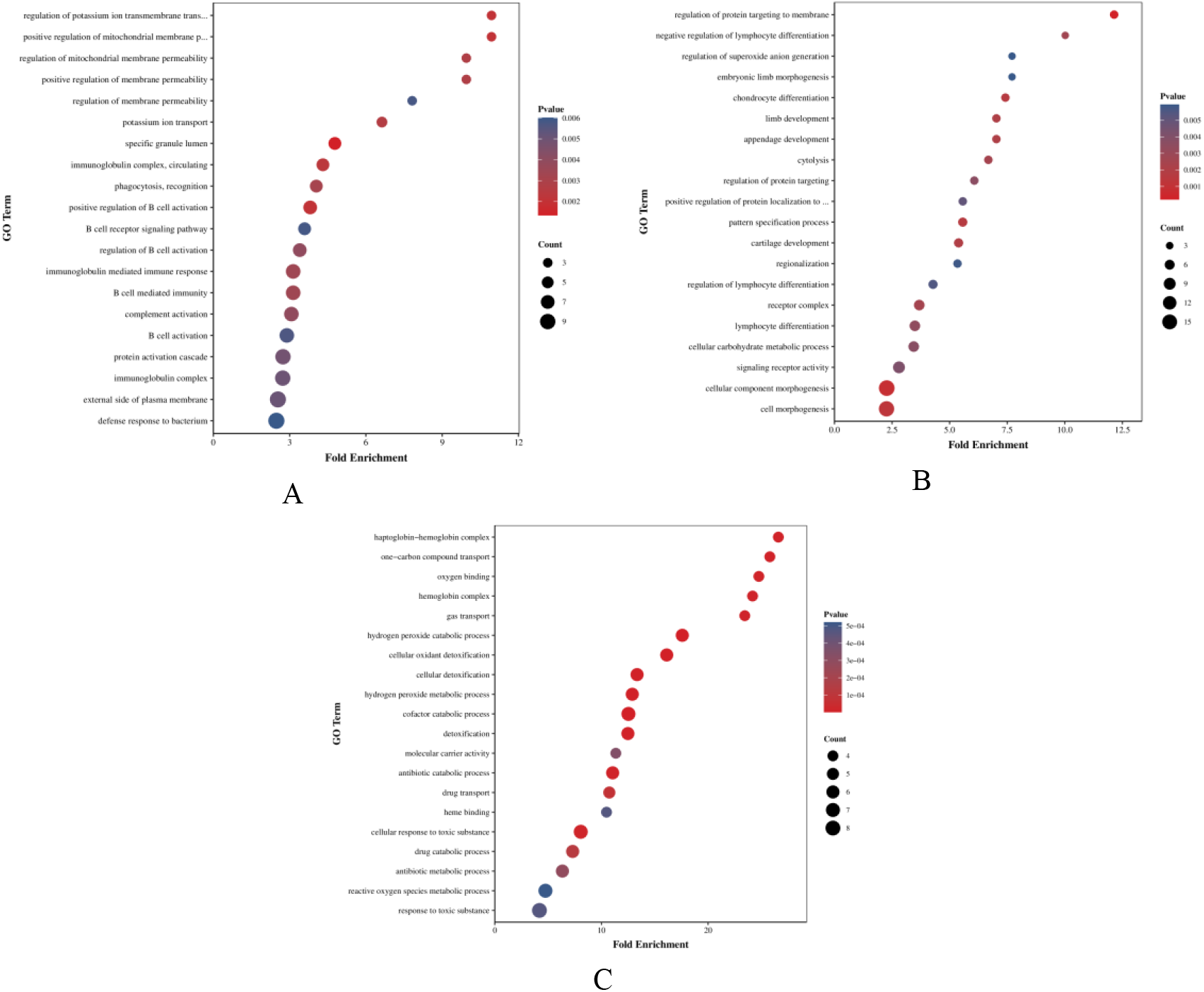
Bubble graph of GO functional enrichment analysis of differentially expressed proteins. Note: A and B respectively represent the top 20 GO-Terms ranked by P-value of differentially expressed proteins in the peripheral serum samples from the low-altitude group and the mid-altitude group at 8d and 28d after chemotherapy; C represents the top 20 GO-Terms ranked by P-value of differentially expressed proteins in the bone marrow supernatant samples from the low-altitude group and the mid-altitude group at 28d after chemotherapy. The horizontal axis represents the enrichment ratio (the ratio of GeneRatio to BgRatio), where a higher enrichment ratio indicates a higher degree of enrichment of differential proteins. The vertical axis represents the names of GO terms. The color of the dots changes from blue to red, representing the change in P-value from large to small, where a smaller P-value indicates greater statistical significance. The size of the dots represents the number of annotated differential proteins for each corresponding term.

The top 20 GO-Terms with enriched analysis results (ranked from smallest to largest P-value) were selected to create an enrichment entry bubble chart. The GO enrichment analysis of differentially expressed proteins in peripheral blood serum at 8 days after chemotherapy shows that these proteins are mainly involved in the regulation of potassium ion transmembrane transport, positive regulation of mitochondrial membrane permeability, regulation of mitochondrial membrane permeability, positive regulation of membrane permeability, membrane permeability regulation, potassium ion transport, specific granule lumen, immunoglobulin complex, circulating, phagocytosis, recognition, and positive regulation of B cell activation. The GO enrichment analysis of differentially expressed proteins in peripheral blood serum at 28 days after chemotherapy shows that these proteins are mainly involved in the regulation of protein targeting to the membrane, negative regulation of lymphocyte differentiation, regulation of superoxide anion generation, embryonic limb morphogenesis, chondrocyte differentiation, limb development, appendage development, cytolysis, regulation of protein targeting, and positive regulation of protein localization to membrane.

The GO enrichment analysis of differentially expressed proteins in bone marrow supernatant at 28 days after chemotherapy shows that these proteins are mainly involved in binding hemoglobin complex, one-carbon compound transport, oxygen-binding, hemoglobin complex, gas transport, hydrogen peroxide catabolic process, cellular oxidant detoxification, cellular detoxification, hydrogen peroxide metabolic process, cofactor catabolic process, and others.

1. The protein counts annotated to all second-level GO terms under the three primary categories were statistically analyzed. The statistical results of GO classification for differentially expressed proteins are shown in the following figure:
  (2) The statistics of the number of proteins annotated to all second-level GO entries under the three primary classifications are shown in the figure below, depicting the GO classification of up-regulated and down-regulated differentially expressed proteins:
2. Select the top 50 GO-Terms ranked by P-value in the enrichment analysis results (sorted from smallest to largest), and plot the enrichment entry bar graph as shown in the figure below:
  (3) Select the top 20 GO-Terms ranked by P-value in the enrichment analysis results (sorted from smallest to largest), and plot the enrichment entry bubble graph as shown in the figure below:

### Enrichment analysis of differentially expressed protein KEGG pathway

In living organisms, different proteins coordinate with each other to carry out their biological functions. Pathway analysis helps to further understand their biological functions. KEGG is a major public database related to pathways (https://www.genome.jp/kegg/). Differential proteins with significant differences are incorporated into the KEGG database for retrieval, resulting in a bar graph showing the enriched distribution of differentially expressed proteins in KEGG pathways, as shown in Figure 8 below.

**Figure 8:**
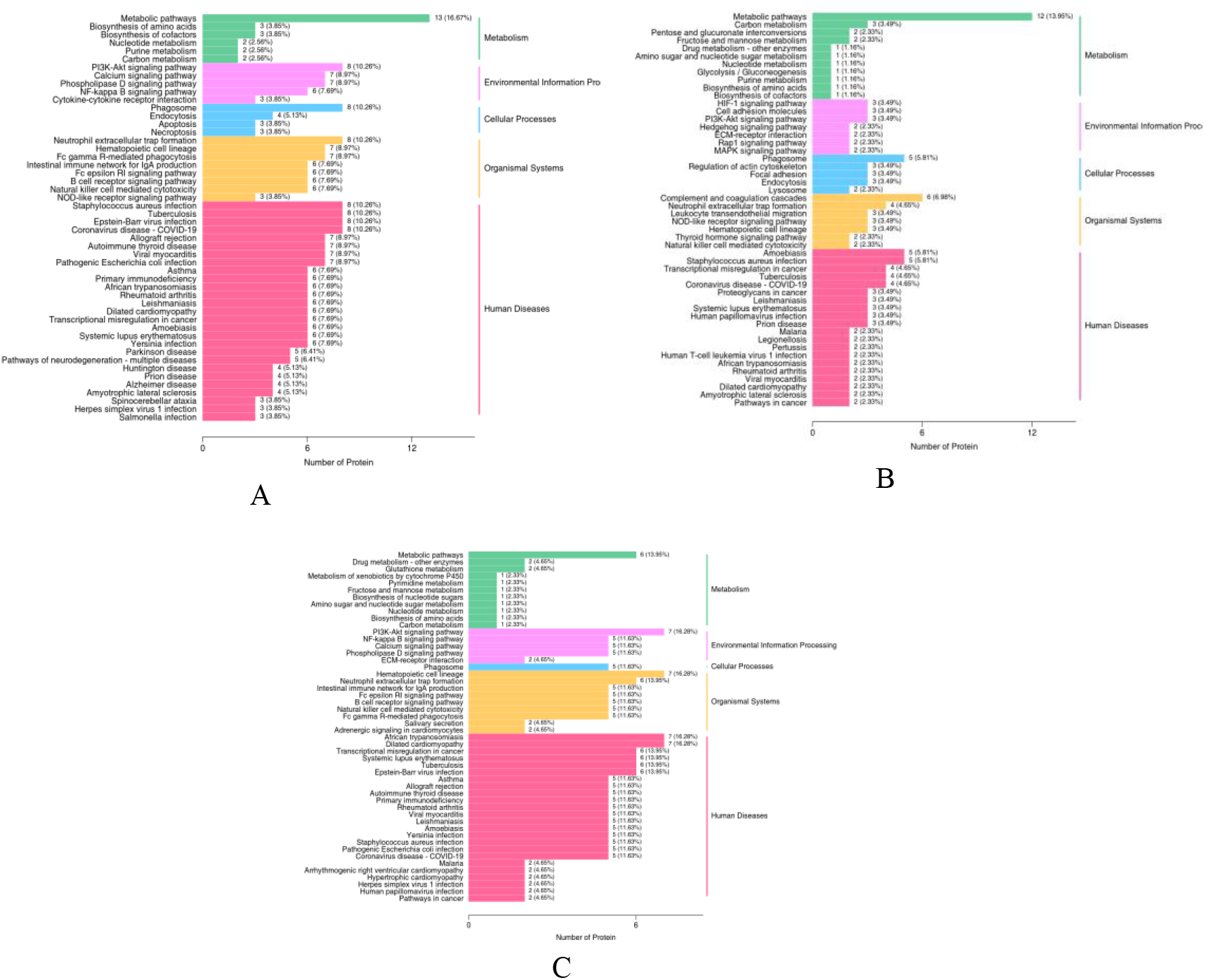
KEGG Classification Bar Graph. Note: A and B respectively represent the KEGG signaling pathways annotated by differential proteins in peripheral serum samples from the low-altitude group and the mid-altitude group at 8d and 28d after chemotherapy; C represents the KEGG signaling pathways annotated by differential proteins in bone marrow supernatant samples from the low-altitude group and the mid-altitude group at 28d after chemotherapy. The horizontal axis represents the number of differential proteins annotated to each pathway, and the vertical axis represents the names of KEGG pathways. The numbers in the graph represent the number of annotated differential proteins for each pathway, with the specific value of the ratio of the number of annotated differential proteins to the total number of annotated differential proteins in parentheses. The label on the far right represents the primary classification to which the KEGG pathway belongs.

**Figure 9:**
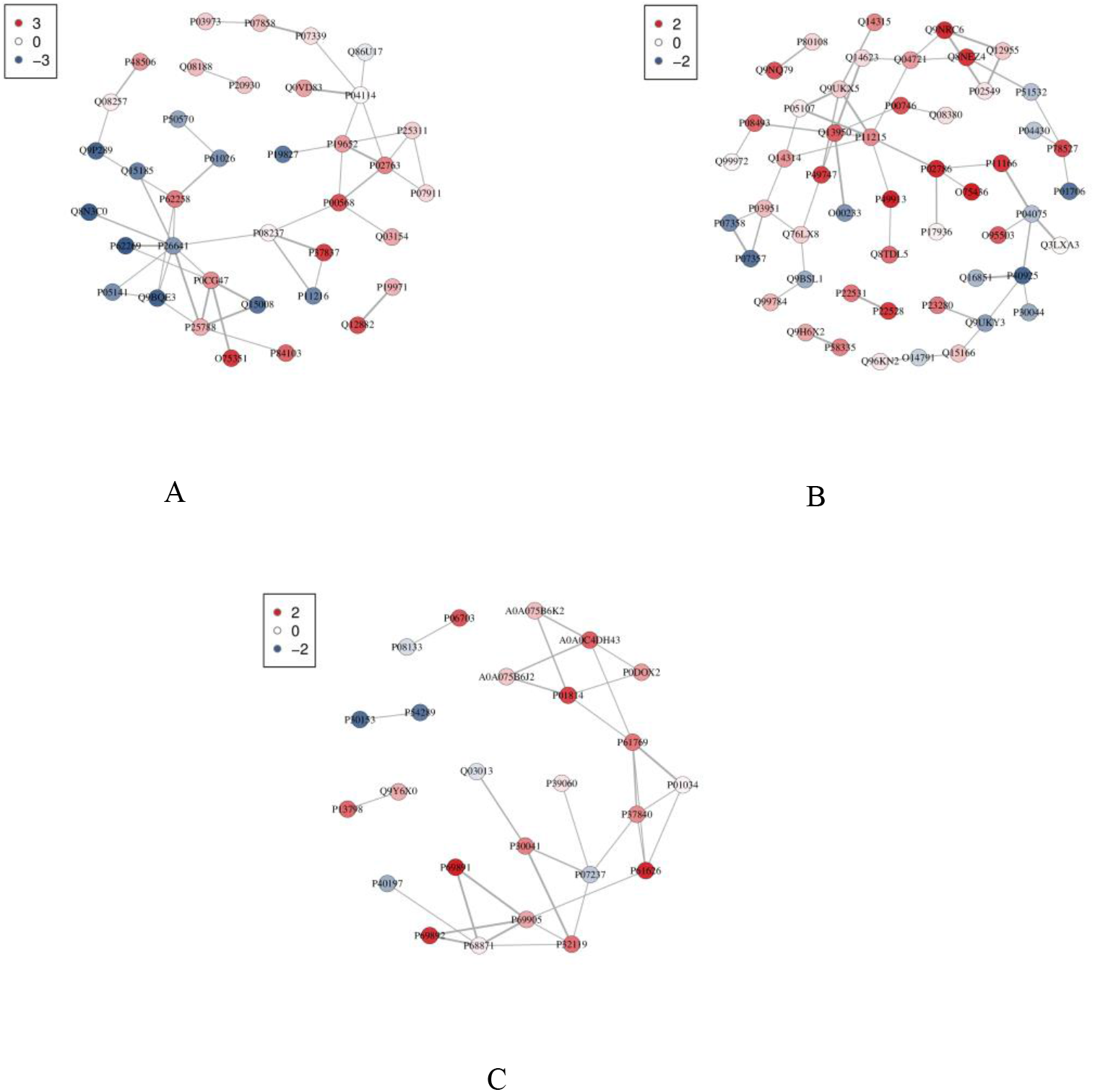
Protein-protein interaction network diagram. Note: A and B represent the protein interaction networks of differentially expressed proteins in the peripheral serum of the low-altitude group and the mid-altitude group at 8d and 28d after chemotherapy, respectively. C represents the protein interaction network of differentially expressed proteins in the bone marrow supernatant of the low-altitude group and the mid-altitude group at 28d after chemotherapy. Each node in the interaction network represents a protein. The color of the nodes ranges from red to blue, indicating changes in protein expression levels from upregulation to downregulation. For differential grouping with 3 or more samples, all nodes are uniformly red. The thickness of the lines represents the change in the credibility of the interaction relationship from high to low.

After performing KEGG pathway enrichment analysis on the differential proteins in peripheral serum samples taken at 8 days post-chemotherapy, the following enrichments were observed: In the metabolism category, significant enrichments were found in metabolic pathways and biosynthesis of amino acids. In the signaling pathways category, enrichments were prominent in the PI3K-Akt signaling pathway, calcium signaling pathway, and phospholipase D signaling pathway. In the cellular processes category, significant enrichments were observed in phagosome, endocytosis, and apoptosis. In the organismal systems category, notable enrichments included neutrophil extracellular trap formation, hematopoietic cell lineage, and Fc gamma R-mediated phagocytosis. In the human diseases category, major enrichments were observed in Staphylococcus aureus infection, tuberculosis, and Epstein-Barr virus infection.

After conducting KEGG pathway enrichment analysis on the differential proteins in peripheral serum samples taken at 28 days post-chemotherapy, the following enrichments were observed: Metabolism category: Significant enrichments were found in metabolic pathways, carbon metabolism, and the interconversion of pentose and glucuronate. Environmental information processing category: Enrichments were prominent in the HIF-1 signaling pathway, cell adhesion molecules, and the PI3K-Akt signaling pathway. Cellular processes category: Significant enrichments were observed in phagosome, and the regulation of the actin cytoskeleton. Organismal systems category: Main enrichments included the complement and coagulation cascades and neutrophil extracellular trap formation. Human diseases category: Major enrichments were observed in amoebiasis and staphylococcus aureus infection.

After conducting KEGG pathway enrichment analysis on the differential proteins in bone marrow supernatant samples taken at 28 days post-chemotherapy, the following enrichments were observed: Metabolism category: Significant enrichments were found in metabolic pathways and drug metabolism-other enzymes. environmental information processing category: Enrichments were prominent in the PI3K-Akt signaling pathway, NF-κB signaling pathway, and calcium signaling pathway. Cellular processes category: Significant enrichments were observed in phagosome. Organismal systems category: Main enrichments included neutrophil extracellular trap formation, hematopoietic cell lineage, and the intestinal immune network for IgA production. Human diseases category: Major enrichments were observed in African trypanosomeiasis, dilated cardiomyopathy, and transcriptional misregulation in cancer.

### Protein Interaction Networks and Modular Analysis

To predict protein-protein interactions, we utilized the StringDB (http://string-db.org/) protein interaction database to construct the PPI network. If the corresponding species were available in the database, we directly extracted the protein sequences of the respective species. If not, we extracted protein sequences from closely related species. The differential protein sequences were then compared with the extracted sequences using blast alignment. Subsequently, interactions of differential proteins were extracted based on a confidence score > 400 (medium confidence). We utilized R packages (graph) and R (networkD3) to construct static and dynamic network graphs respectively. The static graph of the PPI interaction network analysis results is shown below. The corresponding network relationships can be directly imported into Cytoscape software for visualization and editing.

## Discussion

The most common side effect of chemotherapy in AML patients is bone marrow suppression. This is one of the key reasons for dose reduction, treatment limitation, or even discontinuation of therapy(Li *et al*., 2017).After chemotherapy, bone marrow suppression has become a significant threat to the survival rate and quality of life of patients with malignant tumors, necessitating timely and effective preventive and management measures. This also poses a challenging task in the clinical treatment of AML patients. In clinical practice, we have found that even without the influence of physical and age factors, mid-altitude AML patients experience significantly exacerbated bone marrow suppression and prolonged hematopoietic recovery time after standard chemotherapy compared to low-altitude AML patients. Therefore, we conducted proteomic analysis of serum and bone marrow supernatant proteins at 8 days and 28 days after chemotherapy in AML patients at different altitudes. The results showed that at 8 days after chemotherapy, there were 78 differentially expressed proteins between the mid-altitude AML group and the low-altitude AML group in peripheral serum, with 52 upregulated proteins and 26 downregulated proteins. At 28 days after chemotherapy, there were 86 differentially expressed proteins in peripheral serum between the mid-altitude AML group and the low-altitude AML group, with 59 upregulated proteins and 27 downregulated proteins. In addition, at 28 days after chemotherapy, there were 43 differentially expressed proteins in bone marrow supernatant between the mid-altitude AML group and the low-altitude AML group, with 32 upregulated proteins and 11 downregulated proteins. There was an overlap of 1 protein among the three groups, which was Immunoglobulin heavy variable 3-13, showing consistent upregulation and differential trends. Furthermore, there were 2 overlapping differentially expressed proteins between serum samples at 8 days and 28 days after chemotherapy, namely Complement factor D and Fibroleukin. Both proteins showed significantly high expression in both groups, with consistent differential trends, indicating their potential critical role in disease progression.

Subcellular localization of differential proteins indicates that at different time points (8d and 28d after chemotherapy) and different sample sources (peripheral serum and bone marrow supernatant), the subcellular localization of differentially expressed proteins is predominantly in the cytoplasm, extracellular space, and nucleus, followed by mitochondria and membranes, in descending order of percentage. This suggests that the reasons underlying the differences in bone marrow suppression after chemotherapy in low and mid-altitude AML patients may involve several aspects:1. Differences in intracellular environmental regulation: Cytoplasm, nucleus, and mitochondria are important sites for intracellular biological processes. Their different localization may reflect differences in intracellular environmental regulation during bone marrow suppression. These differences may be related to cell survival status, metabolic activity, and stress response factors;2. Differences in extracellular environment: Differential protein localization in the extracellular space also shows some variability, which may reflect the influence of extracellular environments from different sample sources on protein expression and localization. The composition and physiological status of peripheral serum and bone marrow supernatant may differ, thereby affecting the subcellular localization of differential proteins. Changes in metabolism regulation and cellular functions: The cytoplasm and nucleus serve as important centers for cellular metabolism and regulation. Their differential localization may reflect changes in metabolism regulation and cellular functions during bone marrow suppression. These changes may involve important biological processes such as cell signaling transduction and gene transcription regulation. The reasons for the differences in bone marrow suppression after chemotherapy in low and mid-altitude AML patients may involve differences in intracellular and extracellular environmental regulation, as well as related changes in metabolism regulation and cellular functions. Further exploration of these differences will help deepen our understanding of the pathogenesis and treatment response of AML, providing more theoretical basis and clinical guidance for individualized treatment and prognosis assessment. Studies have shown that chemotherapy drugs can affect mitochondrial function by downregulating peroxisome decomposing enzymes, leading to an imbalance between oxidation and antioxidation in the body. This imbalance results in increased reactive oxygen species (ROS) in the matrix cells, causing oxidative damage, inhibiting the proliferation of bone marrow stromal cells (BMSCs), inducing their senescence and apoptosis, and indirectly affecting hematopoietic stem cell (HSC) function(Barjaktarovic *et al*, 2011; Li *et al*, 2020; Xiao *et al*, 2017).ROS (Reactive Oxygen Species) are oxygen-containing compounds generated during aerobic metabolism in cells, and their accumulation can lead to oxidative damage. The ROS produced by normal cells mainly come from the mitochondrial respiratory chain. Damage and dysfunction of mitochondrial complexes can lead to an increase in ROS levels. Liu, Li, and others found that after treatment with the chemotherapy drug daunorubicin (DNR), the bone marrow stromal cell line OP9 cells exhibited extreme mitochondrial expansion, vacuolation of the inner membrane and cristae, decreased mitochondrial membrane potential, and induced production of reactive oxygen species such as superoxide (O2-) and hydrogen peroxide (H2O2). The expression level of the antioxidant enzyme glutathione peroxidase (GPX) decreased, leading to an increase in intracellular ROS levels, triggering oxidative damage to bone marrow stromal cells (BMSCs), an increase in the proportion of senescent cells, a decrease in the ability of normal hematopoietic cell colonies to form, and an increase in DNA damage(Li *et al*., 2020).

Furthermore, research by Xiao et al. found that 5-FU inhibits the colony-forming ability of BMSCs in a concentration-dependent manner. Simultaneously, it leads to the accumulation of ROS in cells, causing cell cycle arrest at the G1 phase, an increase in the positivity rate of senescence-associated β-galactosidase (SA-β-Gal), and an increase in the proportion of apoptotic cells(Xiao *et al*., 2017). In recent years, studies have found that mitochondria in tumor cells can also release excessive ROS. Additionally, an increase in radiation dosage during radiotherapy can lead to enhanced mitochondrial oxidative metabolism(Barjaktarovic *et al*., 2011). The increase in intracellular ROS levels triggers oxidative stress, leading to G1 phase arrest, inhibition of cell proliferation, and modulation of cellular senescence and apoptosis. Thus, mitochondria emerge as crucial organelles regarding the differences in bone marrow suppression after chemotherapy in low and mid-altitude AML patients. GO enrichment analysis indicates that differentially expressed proteins are primarily involved in cellular processes, metabolic processes, and other biological processes. This suggests that protein regulation during bone marrow suppression may influence cellular metabolic activities and other biological processes, which are crucial for bone marrow function recovery.

Metabolic processes are fundamental to cell survival and function, including energy metabolism, substance synthesis, and decomposition. During the bone marrow suppression period after chemotherapy in low and mid-altitude AML patients, the regulation of cellular metabolic activities and the re-adjustment of energy metabolism may lead to differences in bone marrow suppression between low and mid-altitude regions. These changes may be related to cellular adaptation to the external environment. Research has found that metabolic profiling studies of cyclophosphamide-induced bone marrow suppression in mice showed significant changes in plasma metabolites after administration(Chen *et al*, 2017). Nitrophenol is a metabolic product of the aromatic compound nitrobenzene, which can destroy red blood cells and rapidly convert hemoglobin to methemoglobin. This is a relatively stable compound that cannot ionize in tissues and cannot effectively supply oxygen, leading to tissue and organ hypoxia, resulting in decreased red blood cell count and hemoglobin levels. Cellular processes involve various biological processes within the cell, such as cell proliferation, apoptosis, signal transduction, and more. During bone marrow suppression, enrichment of cellular processes may reflect the regulation of cellular responses to external stimuli and damage. Under steady-state conditions, key transcription factors regulate hematopoietic stem cells (HSCs) to maintain them in the G0 phase of the cell cycle. When the body experiences hematologic stress (such as radiation therapy, chemotherapy, and infection) leading to acute bone marrow suppression and depletion of hematopoietic progenitor cells (HPCs), hematopoietic stem cells enter the cell cycle and proliferate and differentiate to replenish HPCs, maintaining homeostasis in the human hematopoietic system. However, toxicity from chemotherapy drugs selective to HSCs, multiple chemotherapy treatments, high-dose radiation therapy, and other factors can impair self-renewal capacity, leading to potential bone marrow damage primarily due to HSC aging. Chemotherapy-induced elevation of ROS, DNA damage, or mitochondrial dysfunction can activate and enhance AMPK, inducing P53-dependent cell cycle arrest. Additionally, DNA damage can recruit ATM/ATR to activate P53, ultimately leading to cell cycle arrest at the G1 phase through the activation of retinoblastoma protein (Rb), thereby inducing HSC aging(Sorimachi *et al*, 2021). The immune response is a crucial mechanism by which the body combats invading pathogens and foreign substances. During bone marrow suppression, the regulation of the immune response may involve processes such as the release of inflammatory factors, activation, and regulation of immune cells, among others. These changes may be related to the restoration of patient immune function and the modulation of infection risk. This also suggests that there may be differences in the response to infections during the bone marrow suppression period after chemotherapy in low and mid-altitude AML patients.

The KEGG pathway enrichment analysis results indicate that the occurrence of bone marrow suppression differences after chemotherapy in low and mid-altitude AML patients is regulated through the coordinated action of multiple pathways rather than being summarized by a single pathway. Currently, our focus lies on the PI3K-Akt signaling pathway, HIF-1 signaling pathway, NF-κB signaling pathway, and calcium signaling pathway.

The PI3K/AKT signaling pathway is a pathway for transducing signals that respond to extracellular survival and growth signals in cells. It serves as an important mediator in the regulation of hematopoietic function through cytokine signal transduction. Inhibiting the PI3K-AKT signaling pathway can improve the hematopoietic differentiation of embryonic stem cells and promote their differentiation into hematopoietic progenitor cells (HPCs) (Nii *et al*, 2015). However, excessive activation of the PI3K-AKT pathway may lead to depletion of hematopoietic stem cells (Kharas *et al*, 2010). PI3K is composed of a regulatory subunit (P85) and a catalytic subunit (P110). The P85 subunit is associated with the tyrosine residues (Y607 and Y508 sites) of EPO-R, activating the PI3K pathway. Activated PI3K converts PIP2 on the cell membrane to PIP3. PIP3 activates PDK1, leading to its phosphorylation and activation of Akt (protein kinase B). Akt is considered the central mediator of PI3K/AKT signal transduction. Activated Akt can promote cell growth and proliferation through multiple mechanisms, including inhibiting apoptosis, promoting protein synthesis, and regulating the cell cycle. Akt nuclear translocation is necessary for EPO-induced erythroid differentiation. Subsequently, downstream targets of PI3K/AKT are activated, including transcription factors FOXO3 and GATA-1, hypoxia-inducible factor-1α (HIF1α), and mammalian targets of rapamycin (mTOR), which are critical for normal red blood cell development(Zhang *et al*, 2018). Additionally, Akt kinase activation can promote cell cycle progression and inhibit apoptosis, thereby maintaining the quantity and function of hematopoietic stem cells (HSCs), exerting a significant impact on HSC proliferation and differentiation(Adlung *et al*, 2017; Mei *et al*, 2021; Tóthová *et al*, 2021).Zhang et al. found that Compound Donkey-Hide Gelatin Syrup could promote the recovery of 5-fluorouracil-induced bone marrow suppression by modulating the PI3K signaling pathway(Zhang *et al*, 2019). Furthermore, we found that differentially expressed proteins in serum at 8 and 28 days after chemotherapy and in bone marrow supernatant at 28 days after chemotherapy are enriched in the PI3K/AKT signaling pathway. Therefore, the regulation of the PI3K/AKT pathway is closely associated with the differences in bone marrow suppression after chemotherapy in low and mid-altitude groups.

The low oxygen environment in the microenvironment is often a significant pathological characteristic of inflammatory tissues. Hypoxia is frequently accompanied by the infiltration of inflammatory cells, and cellular hypoxic responses are closely intertwined with the body’s inflammatory reactions(Gilreath *et al*, 2014).The HIF family consists of α and β subunits, with HIF-1α being the primary active component of HIF-1. Under normal oxygen conditions, HIF-1α is recognized and hydroxylated by the PHD family. Hydroxylated HIF-1α binds to the von Hippel-Lindau tumor suppressor protein (pVHL), forming an E3 ubiquitin ligase complex, which ultimately degrades HIF-1α through the ubiquitin-proteasome pathway. The prolyl hydroxylases (PHDs) in the non-heme iron-dependent dioxygenase family hydroxylate HIF-1α in the presence of oxygen. Under hypoxic conditions, PHD hydroxylation activity decreases, inhibiting the oxygen-dependent degradation pathway of HIF-1α. This stabilizes HIF-1α within the cell, allowing it to bind with HIF-1β to form the active HIF-1 complex.

This complex binds to hypoxia response elements (HREs) and participates in various biological processes, including inflammation and immune regulation(Wang *et al*, 2021). Due to factors such as oxygen concentration and immune response metabolites, among others, participating in the regulation of HIF-1α activation and the inflammatory response process(Luo *et al*, 2017), we believe that the abnormal metabolic microenvironment caused by post-chemotherapy bone marrow suppression, along with the involvement of altitude factors through this metabolic pathway, leads to differences in the degree of post-chemotherapy bone marrow suppression in AML patients at different altitudes. Nuclear factor-kappa B (NF-κB) is a class of nuclear transcription factors that can specifically bind to the enhancer κB sequence of the immunoglobulin κ light chain gene. Research has confirmed increased activation of NF-κB in aging hematopoietic stem cells (HSCs). Studies have found that NF-κB and ATM are co-activated in cells and tissues with higher levels of DNA damage-induced aging. DNA damage triggers the activation of the PIDDosome (containing RIP1/PIDD/RAIDD), leading to the sulfhydration of NEMO and its translocation to the nucleus. Simultaneously, DNA damage activates ATM, which together with sulfhydrated NEMO, induces NEMO phosphorylation and ubiquitination, leading to the return of the cytoplasm and mediating IKK activation. This process subsequently phosphorylates IκB protein and dissociates it from the trimer, promoting the transcription of related genes, mediating hematopoietic stem cell aging, apoptosis, and the production of SASP(Hayden *et al*, 2006; Martínez-Zamudio *et al*, 2017).In 2018, Chen et al. proposed that the increased activity of NF-κB during the basal phase of inflammation in aged mice HSCs and the failure to downregulate this pathway during inflammation resolution are closely related to HSC aging. They also indicated that Rad21/cohesin plays a significant role in mediating NF-κB signaling in HSPCs (Rad21/cohesin deficiency has little effect on gene regulation under steady-state conditions). Moreover, the activation of the NF-κB signal has a stronger impact on the self-renewal capacity of aged mouse HSCs(Chen *et al*, 2019). Additionally, NF-κB is involved in inflammaging. A study by He et al. published in *Blood* in 2021 demonstrated that elevated pro-inflammatory cytokine TNF-α in the bone marrow can increase IL27Ra transcriptional activity by activating ETS1 and NF-κB p65, thereby promoting HSC aging(He *et al*, 2020).Furthermore, the subunit p65 of NF-κB shares common binding sites with the promoter of HIF-1a. NF-κB can enhance the regulatory function of HIF-1a in inflammatory responses. In the absence of NF-κB, even under prolonged exposure to hypoxic conditions, the HIF-1a gene cannot be effectively transcribed. However, HIF-1a can mediate the activation of NF-κB in hypoxic cells, promoting the regulation of cytokine expression in macrophages by NF-κB. This implies that HIF-1a and NF-κB interact in immune responses, forming the HIF-1a/NF-κB signaling pathway, which regulates immune cell activity and promotes the expression of pro-inflammatory factors(Choi *et al*, 2014; Görlach & Bonello, 2008).

In summary, this study investigated the impact of altitude on protein expression during the bone marrow suppression period after chemotherapy in patients with low and intermediate altitude AML. By analyzing the proteins in serum and bone marrow supernatant post-chemotherapy and employing bioinformatics analysis, the study aimed to explore the influence of altitude on protein expression in AML patients during bone marrow suppression. The results revealed differences in the proteomics profiles between the two groups of patients during bone marrow suppression post-chemotherapy, with the differentially expressed proteins primarily involved in cellular and metabolic processes among other biological pathways. Further analysis indicated that these differential proteins exert their effects through the PI3K-Akt, HIF-1, NF-κB, and calcium signaling pathways. These findings provide valuable insights into the differences in bone marrow suppression severity after chemotherapy at different altitudes. However, as this study was based on proteomics and bioinformatics analyses, further experimental validation is needed to elucidate the mechanisms by which altitude regulates these signaling pathways.

## Materials and Methods

### Data Collection

Between 2021 and 2022, a total of 5 newly diagnosed and untreated AML patients (non-M3 subtype) were selected from the Hematology Departments of Qinghai Provincial People’s Hospital (located at a moderate altitude of 1,501-2,500 meters) and the Xi Jing Hospital of the Fourth Military Medical University (located at a low altitude of 500-1,500 meters). After obtaining approval from the Ethics Committee of Qinghai Provincial People’s Hospital (Ethics Approval No: 2022-25) and informed consent from the participants, general information of the participants was collected.

Inclusion Criteria:

1. Diagnosis of AML based on the latest MICM criteria (morphology, immunology, cytogenetics, and molecular biology) with FAB classification.
2. Received standard induction chemotherapy regimen: cytarabine at a dose of 100-200 mg/m^2^/day for 7 days combined with daunorubicin at a dose of 12 mg/m^2^/day for 3 days, or idarubicin at a dose of 60-90 mg/m^2^/day for 3 days.

Exclusion Criteria:

1. AML related to previous treatment.
2. Patients with severe acute or chronic infectious diseases.
3. Patients with contraindications to chemotherapy or those who did not complete one cycle of chemotherapy.

These criteria were applied to ensure the consistency and integrity of the study population. Marrow suppression grading: Marrow suppression grading for patients at days 8, 14, and 28 after chemotherapy was determined using the WHO criteria for acute and subacute toxicity reactions to anticancer drugs.

Blood and marrow samples (2 ml each) were collected before chemotherapy and on days 8 and 28 after standard initial chemotherapy (peripheral blood only) using serum biochemical tubes. Samples were centrifuged at 1600 g for 10 minutes at 4°C to obtain supernatants. The supernatants were then subjected to high-speed centrifugation at 12000 r for 2 minutes at 4°C, and the upper layer of serum was transferred to sterile, enzyme-free storage tubes (0.1 ml per tube) and stored at -80°C. Protein profiling analysis was performed using 4D-DIA.

### DIA protein quantification

#### Main reagents

Bovine Serum Albumin (Wuhan Chucheng Zhengmao Technology Engineering Co., Ltd.), Dithiothreitol (Solarbio), Ethylenediaminetetraacetic Acid (EDTA) (National Medicines), Coomassie Brilliant Blue G-250 (National Medicines), Iodoacetamide (Aladdin), N-Benzoyl-Sulfonyl Fluoride (Xiya Reagent), Sodium Dodecyl Sulfate (SDS) (National Medicines), Tetraethylammonium Bromide (Sigma), Thiourea (National Medicines), Tris(hydroxymethyl)aminomethane (Solarbio), Trypsin (Promega), Urea (Sigma), Protein Marker (Fermentas), Acetone (National Medicines), BCA Protein Quantification Kit (Beyotime).

#### Sample preparation and proteolysis

1. Follow the instructions of the ProteoMiner™ Protein Enrichment Small-Capacity Kit (1633006, Bio-Rad) to remove high-abundance proteins from the samples and collect the final wash eluate.
2. Add 4 times the volume of cold acetone and 10 mM DTT to the samples, precipitate at -20°C for 2 hours, centrifuge at 13,000 g for 20 minutes at 4°C, discard the supernatant, air dry the protein precipitate, and resuspend the protein in 8M urea/100mM TEAB (pH 8.0).
3. Add DTT to a final concentration of 10 mM and perform a reduction reaction at 56°C for 30 minutes. Add IAM to a final concentration of 55 mM and allow it to react in the dark at room temperature for 30 minutes for alkylation.
4. Add 4 times the volume of cold acetone and 10 mM DTT, precipitate at -20°C for 2 hours, centrifuge at 13,000 g for 20 minutes at 4°C, discard the supernatant.
5. Resuspend the protein in 8M urea/100mM TEAB (pH 8.0) solution.
6. Determine the protein concentration using the BCA method. Divide the samples and store at - 80°C.
7. Take an appropriate amount of protein for 12% SDS-PAGE quality control. (Note: Typically, 30 μg of protein is used for Coomassie staining, and 2 μg of protein is used for silver staining for quality control).
8. Desalt the enzymatic digestion. Take 100 μg of protein from each sample for trypsin digestion. Dilute the protein solution 5 times with 100 mM TEAB, then add trypsin at a mass ratio of 1:50 (trypsin:protein). Digest overnight at 37°C. Use a C18 Cartridge to desalt the digested peptides, and vacuum freeze-dry the desalted peptides. The freeze-dried peptide powder is resuspended in 0.1% formic acid water to a concentration of 0.1 μg/μl and stored at -20°C for later use.

#### LC-MS/MS detection

##### 1. Nano Elut high-performance liquid chromatograph

The samples were separated using the NanoElute system with a flow rate of nanoliters. The mobile phase consisted of 0.1% formic acid aqueous solution (Phase A) and 0.1% formic acid acetonitrile aqueous solution (acetonitrile is 100% B).

The samples were automatically injected into the analytical column (IonOpticks, Australia, 25cm X 75μm, C18, 1.6μm) for separation. The column temperature was controlled at 50°C by the integrated column oven. The injection volume was 200 ng, the flow rate was 300 nL/min, and the gradient was 60 minutes. The liquid phase gradient over 60 minutes was as follows:

- 0 min to 45 min: Phase B increased from 2% to 22%
- 45 min to 50 min: Phase B increased linearly from 22% to 35%
- 50 min to 55 min: Phase B increased linearly from 35% to 80%
- 55 min to 60 min: Phase B was maintained at 80%.

##### 2. timsTOF Pro2 mass spectrometer

First, the mixed samples were subjected to chromatographic separation, followed by mass spectrometry data acquisition using the ddaPASEF mode of the timsTOF Pro2 mass spectrometer to establish appropriate acquisition windows for the diaPASEF method. The analysis was conducted with an effective gradient of 60 minutes. The detection mode was positive ionization, with a precursor ion scan range of 100-1700 m/z. The ion mobility range was 0.7-1.4 Vs/cm^2, with an ion accumulation and release time of 100 ms. The ion utilization rate was nearly 100%. The capillary voltage was set at 1400V, the drying gas flow rate was 3 L/min, and the drying temperature was 180°C.

Parameters for DDA-PASEF acquisition mode were as follows: 10 MS/MS scans (total cycle time of 1.17s), charge range of 0-5, dynamic exclusion time of 0.4 min, ion target intensity of 10000, ion intensity threshold of 2500, CID fragmentation energy of 42 eV. Isolation window settings: 2 Th for less than 700Th, and 3 Th for greater than 700Th.

Parameters for diaPASEF acquisition mode were as follows: Mass Range approximately 400-1200, Mobility Range 0.7-1.4 V⋅s/cm^2, Mass Width 25Da, Mass Overlap 0.1, Mass Steps per Cycle 32, Number of Mobility Windows 2, totaling 64 acquisition windows. The average acquisition cycle was 1.8s.

#### Database searches

In this study, DIA-NN (version 1.8.1) was used as the database search software, employing the Library-free method. The search parameters were set as follows: The database used was swissprot_Homo_sapiens_9606_20376.fasta database, containing a total of 20376 sequences. The option for deep learning-based parameter prediction of a spectral library was selected. The MBR option was checked to utilize DIA data to generate a spectral library, and this library was used for the reanalysis of DIA data to obtain protein quantification. Both precursor ion and protein-level FDR were filtered at 1%. The filtered data was then available for subsequent bioinformatics analysis.

#### Bioinformatics analysis

The differentially expressed proteins were identified using the DIA proteomics method. In the low and mid-altitude groups, proteins with a fold change (FC) > 1.5 or < 0.6667 and a p-value < 0.05 were defined as significantly differentially expressed proteins. GO and KEGG databases were utilized for bioinformatics analysis of the differentially expressed proteins. The String database was employed to analyze the selected differentially expressed proteins and construct a protein-protein interaction (PPI) network.

#### Statistical analysis

The data results were organized and analyzed using the statistical software SPSS 25.0. Continuous data were presented as mean ± standard deviation. The Mann-Whitney U test was used for comparing two independent samples with skewed distributions, while the Wilcoxon signed-rank test was employed for comparing two related samples with skewed distributions. The significance level was set at α = 0.05 for all tests.

## Data Availability Statement

The data generated in this study are available upon request from the corresponding author.

## Acknowledgments

Thanks to the assistance and support provided by the Air Force Medical University of the Chinese People’s Liberation Army in this study. Thanks to all the volunteers who provided blood and bone marrow samples for this study. Thank you to the following organizations for providing financial support for this article:1. National Blood System Disease Clinical Medical Research Center Branch (2021WWB04). 2.Qinghai Province Blood System Disease Clinical Medical Research Center (2021-SF-136).

## Author contribution statement

Author contributions: Qi Sun proposed the main research objectives, was responsible for the conception and design of the study, implementation of the research, and writing of the manuscript; Qi Sun, Houfa Zhou, and Aibo Wang collected specimens, gathered and organized data, performed statistical analysis, and prepared tables; Wenqian Li and Youbang Xie revised the manuscript; Wenqian Li was responsible for quality control and review of the article, overall responsible for the article, and supervised the management. Thanks to the Department of Hematology, Xijing Hospital, Air Force Medical University of the Chinese People’s Liberation Army for their assistance and support in this study.

## References

Adlung L, Kar S, Wagner MC, She B, Chakraborty S, Bao J, Lattermann S, Boerries M, Busch H, Wuchter P et al (2017) Protein abundance of AKT and ERK pathway components governs cell type-specific regulation of proliferation. Mol Syst Biol 13: 904

Barjaktarovic Z, Schmaltz D, Shyla A, Azimzadeh O, Schulz S, Haagen J, Dörr W, Sarioglu H, Schäfer A, Atkinson MJ et al (2011) Radiation-induced signaling results in mitochondrial impairment in mouse heart at 4 weeks after exposure to X-rays. PLoS One 6: e27811

Chen J, Liu Y, Xu Y, Wang Y (2017) [Analysis on metabolites with small molecule of serum in bone marrow suppression model mice with metabolomics method]. Se Pu 35: 1312–1316

Chen Z, Amro EM, Becker F, Hölzer M, Rasa SMM, Njeru SN, Han B, Di Sanzo S, Chen Y, Tang D et al (2019) Cohesin-mediated NF-κB signaling limits hematopoietic stem cell self-renewal in aging and inflammation. J Exp Med 216: 152–175

Choi JK, Kim KH, Park SR, Choi BH (2014) Granulocyte macrophage colony-stimulating factor shows anti-apoptotic activity via the PI3K-NF-κB-HIF-1α-survivin pathway in mouse neural progenitor cells. Mol Neurobiol 49: 724–733

Gilreath JA, Stenehjem DD, Rodgers GM (2014) Diagnosis and treatment of cancer-related anemia. Am J Hematol 89: 203–212

Görlach A, Bonello S (2008) The cross-talk between NF-kappaB and HIF-1: further evidence for a significant liaison. Biochem J 412: e17–19

Hayden MS, West AP, Ghosh S (2006) SnapShot: NF-kappaB signaling pathways. Cell 127: 1286–1287

He H, Xu P, Zhang X, Liao M, Dong Q, Cong T, Tang B, Yang X, Ye M, Chang Y et al (2020) Aging-induced IL27Ra signaling impairs hematopoietic stem cells. Blood 136: 183–198

Kharas MG, Okabe R, Ganis JJ, Gozo M, Khandan T, Paktinat M, Gilliland DG, Gritsman K (2010) Constitutively active AKT depletes hematopoietic stem cells and induces leukemia in mice. Blood 115: 1406–1415

Li X, Wang W, Chen J (2017) Recent progress in mass spectrometry proteomics for biomedical research. Sci China Life Sci 60: 1093–1113

Li Y, Xue Z, Dong X, Liu Q, Liu Z, Li H, Xing H, Xu Y, Tang K, Tian Z et al (2020) Mitochondrial dysfunction and oxidative stress in bone marrow stromal cells induced by daunorubicin leads to DNA damage in hematopoietic cells. Free Radic Biol Med 146: 211–221

Luo F, Zou Z, Liu X, Ling M, Wang Q, Wang Q, Lu L, Shi L, Liu Y, Liu Q et al (2017) Enhanced glycolysis, regulated by HIF-1α via MCT-4, promotes inflammation in arsenite-induced carcinogenesis. Carcinogenesis 38: 615–626

Martínez-Zamudio RI, Robinson L, Roux PF, Bischof O (2017) SnapShot: Cellular Senescence Pathways. Cell 170: 816-816.e811

Mei Y, Liu Y, Ji P (2021) Understanding terminal erythropoiesis: An update on chromatin condensation, enucleation, and reticulocyte maturation. Blood Rev 46: 100740

Nian J, Sun X, Guo J, Yan C, Wang X, Yang G, Yang L, Yu M, Zhang G (2016) Efficacy and safety of acupuncture for chemotherapy-induced leucopoenia: protocol for a systematic review. BMJ Open 6: e010787

Nii T, Marumoto T, Kohara H, Yamaguchi S, Kawano H, Sasaki E, Kametani Y, Tani K (2015) Improved hematopoietic differentiation of primate embryonic stem cells by inhibition of the PI3K-AKT pathway under defined conditions. Exp Hematol 43: 901-911.e904

Sorimachi Y, Karigane D, Ootomo Y, Kobayashi H, Morikawa T, Otsu K, Kubota Y, Okamoto S, Goda N, Takubo K (2021) p38α plays differential roles in hematopoietic stem cell activity dependent on aging contexts. J Biol Chem 296: 100563

Tóthová Z, Šemeláková M, Solárová Z, Tomc J, Debeljak N, Solár P (2021) The Role of PI3K/AKT and MAPK Signaling Pathways in Erythropoietin Signalization. Int J Mol Sci 22

Wang P, Ding J, Yang G, Sun W, Guo H, Zhao Y (2021) Study on the Mechanism of Qigu Capsule in Upregulating NF-κB/HIF-1α Pathway to Improve the Quality of Bone Callus in Mice at Different Stages of Osteoporotic Fracture Healing. Evid Based Complement Alternat Med 2021: 9943692

Xiao H, Xiong L, Song X, Jin P, Chen L, Chen X, Yao H, Wang Y, Wang L (2017) Angelica sinensis Polysaccharides Ameliorate Stress-Induced Premature Senescence of Hematopoietic Cell via Protecting Bone Marrow Stromal Cells from Oxidative Injuries Caused by 5-Fluorouracil. Int J Mol Sci 18

Zhang Y, Ye T, Hong Z, Gong S, Zhou X, Liu H, Qian J, Qu H (2019) Pharmacological and transcriptome profiling analyses of Fufang E’jiao Jiang during chemotherapy-induced myelosuppression in mice. J Ethnopharmacol 238: 111869

Zhang Z, Yao L, Yang J, Wang Z, Du G (2018) PI3K/Akt and HIF-1 signaling pathway in hypoxia-ischemia (Review). Mol Med Rep 18: 3547–3554

